# Material Properties of Nonpregnant and Pregnant Human Uterine Layers

**DOI:** 10.1101/2023.08.07.551726

**Authors:** Daniella M. Fodera, Serena R. Russell, Johanna L. Lund-Jackson, Shuyang Fang, Xiaowei Chen, Joy-Sarah Y. Vink, Michelle L. Oyen, Kristin M. Myers

## Abstract

The uterus has critical biomechanical functions in pregnancy and undergoes dramatic material growth and remodeling from implantation to parturition. The intrinsic material properties of the human uterus and how they evolve in pregnancy are poorly understood. To address this knowledge gap and assess the heterogeneity of these tissues, the time-dependent material properties of all human uterine layers were measured with nanoindentation. The endometrium-decidua layer was found to be the least stiff, most viscous, and least permeable layer of the human uterus in nonpregnant and third-trimester pregnant tissues. In pregnancy, endometrium-decidua becomes stiffer and less viscous with no material property changes observed in the myometrium or perimetrium. Additionally, uterine material properties did not significantly differ between third-trimester pregnant tissues with and without placenta accreta. The foundational data generated by this study will facilitate the development of physiologically accurate models of the human uterus to investigate gynecologic and obstetric disorders.

**Highlights:**

- Human uterine layers are distinct, heterogeneous and time-dependent
- Pregnancy alters the material properties of the maternal-fetal interface
- Largest dataset of uterine mechanical properties measured by nanoindentation

**Graphical Abstract:** 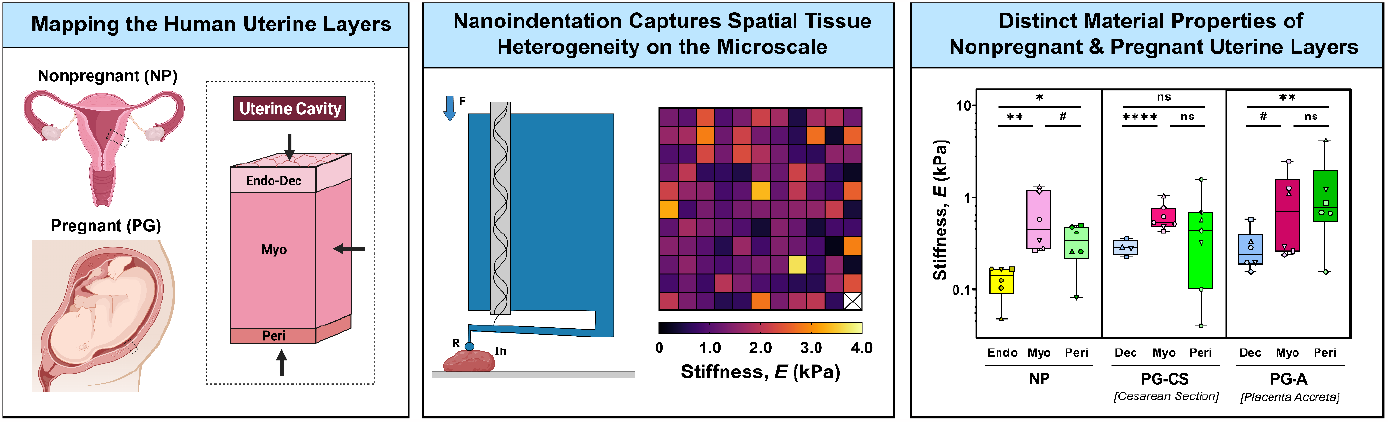

## 1. Introduction

Each year, over 200 million women become pregnant worldwide, with millions more struggling with infertility^1–3^. Even after implantation occurs and a pregnancy is established, complications can arise at every gestational stage, accounting for 20% of all pregnancies^4,5^. Preterm labor, preeclampsia, intrauterine growth restriction, and placenta accreta are amongst the multitude of obstetric conditions that can cause severe morbidity and mortality for both mother and fetus^6–11^. The uterus is integral to the establishment and maintenance of pregnancy, and possesses key biomechanical functions, undergoing extensive growth and remodeling to support fetal development^11^. Defects to mechanical properties of this organ are thought to contribute to the pathogenesis of many obstetric disorders, yet, this remains an open question^6,8,10,11^.

The uterus is composed of three layers (Fig. 1A): (i) the endometrium-decidua, (ii) the myometrium, and (iii) the perimetrium^11–13^. Each of these layers is structurally distinct, though together, they function to support basic reproductive processes. The endometrium is the innermost mucosal layer of the nonpregnant uterus and is composed of epithelial glands and stromal cells embedded in a collagenous matrix lined on its surface with a layer of luminal epithelium^11,14^. Under the influence of hormones, the endometrium remodels dramatically throughout the menstrual cycle^11^. During the proliferative phase, the endometrium rapidly regenerates and thickens, undergoing further differentiation (i.e., decidualization) during the secretory phase^11,15,16^. In the absence of an implanting embryo, menstruation occurs, resulting in endometrial shedding (*4*). Adjacent to the endometrium is the myometrium, the thickest uterine layer composed primarily of smooth muscle fascicles sheathed with an extracellular matrix of collagen and elastic fibers^11,17,18^. Even in its nonpregnant state, the myometrium undergoes active peristaltic contractions throughout the menstrual cycle to aid in sperm motility, embryo implantation, and menstrual blood egress^13,19,20^. Lastly, the perimetrium, also known as the serosa, is the outermost layer of the uterus composed of collagen embedded with epithelial cells that secrete lubricating fluids^12,13,21^.

**Fig. 1.**
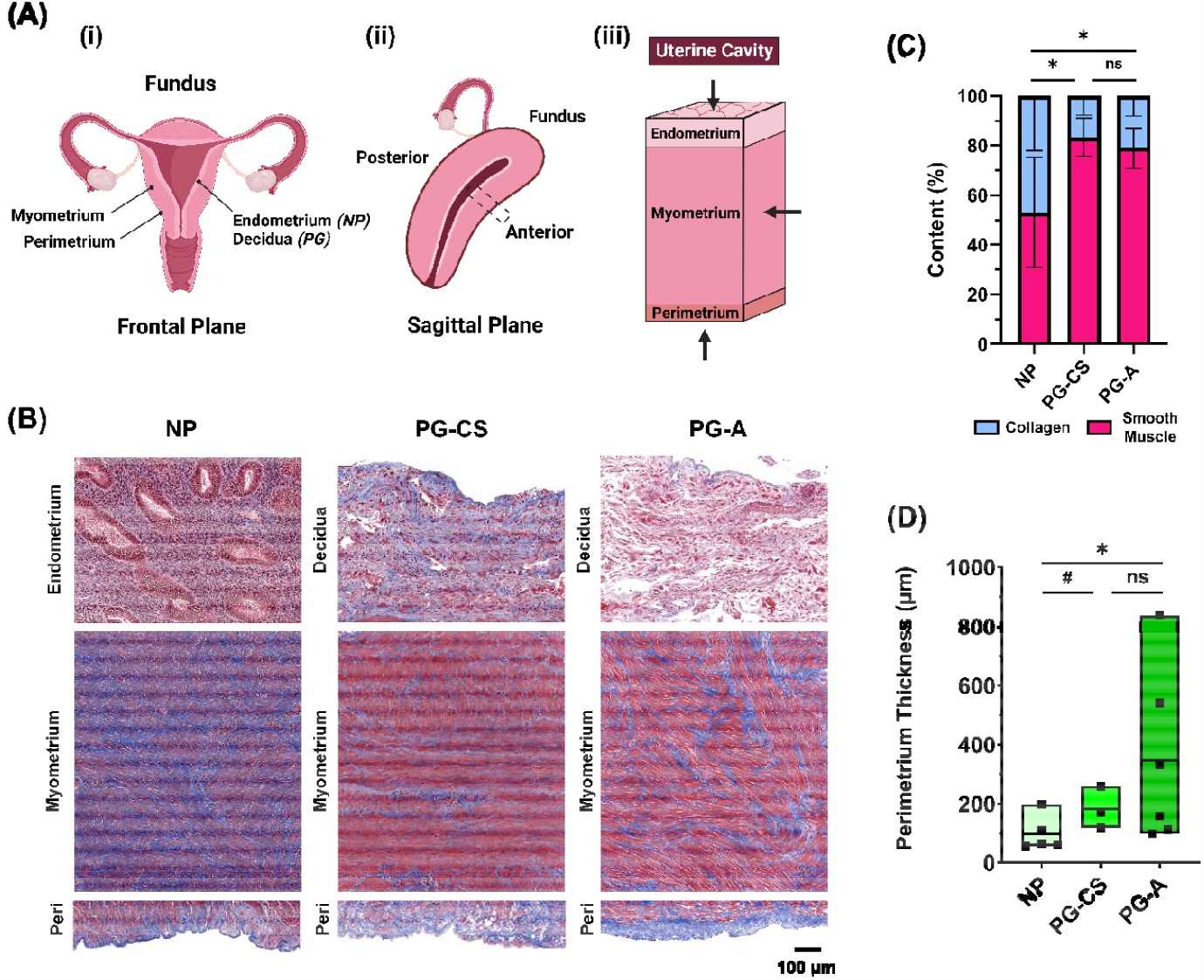
Anatomy of the Human Uterus. **(A)** Diagram of the (i) female reproductive tract in the frontal plane identifying all uterine layers: NP endometrium, PG decidua, myometrium, and perimetrium, (ii) human uterus in the sagittal plane indicating the anterior region from which samples were collected, (iii) cross-section of the uterine wall containing three uterine layers. Arrows indicate the orientation in which tissues were tested with nanoindentation. **(B)** Representative images of the three uterine layers from NP, cesarean section pregnancies (PG-CS), and placenta accreta pregnancies (PG-A) patient groups stained with Masson’s Trichrome (blue = collagen; red = muscle fibers, cytoplasm; black = nuclei). Images were taken at 10x magnification (scale bar = 100 µm). Note that the relative lengths of the tissue layer figure panels do not reflect actual layer proportions. **(C)** Relative proportion of collagen and smooth muscle in the myometrium. Bars indicate standard deviation. **(D)** Thickness of the perimetrium measured for all three patient groups. Data presented as min-max plots. Statistical significance is denoted by the following symbols: ns, p > 0.05; # p _≤_ 0.1; ^*^ p ≤ 0.05.

Each of these distinct uterine layers function in concert to enable and support pregnancy. Following fertilization, implantation of an embryo into the decidualized endometrium is necessary for establishing a successful pregnancy and must occur during a short window in the secretory phase of the menstrual cycle^11,22^. In the process of implantation, trophoblast cells invade through the luminal endometrial epithelium, remodeling the maternal spiral arteries, and form the basis of the placenta^23,24^. The decidua, alongside the placenta, acts as a critical interface between the mother and fetus, which continues to remodel throughout pregnancy under the influence of hormones, to support fetal growth^11,16,25^. Invasive implantation beyond the endometrium-decidua layer into the myometrium, and, in severe cases, through the perimetrium is known as placenta accreta spectrum disorder^9,26,27^. This obstetric condition, which occurs in 0.3% of pregnancies, can lead to severe maternal hemorrhaging, permanent loss of fertility, and mortality for both mother and fetus^9,26,28^. Any prior uterine surgery, including a cesarean delivery, is a major risk factor for placenta accreta, yet the pathophysiology of this disease is poorly understood^29,30^. It is presently unknown whether abnormally invasive placentation is caused by an anomaly in trophoblast biology or the presence of cesarean scar tissue^28,31–33^. Further, the potential contribution of mechanics in the initiation and progression of placenta accreta has yet to be investigated.

In normal pregnancy, the myometrium also experiences dramatic growth and remodeling, exhibiting a nearly twenty-fold increase in volume, to accommodate the growing fetus^11,13,34^. This expansion is initially achieved through hyperplasia, an increase in cell number, in the first trimester, followed by stretch-induced hypertrophy, an increase in cell size, during the later stages of pregnancy^34^. The dynamic activity of the myometrium peaks at the time of labor in the form of coordinated and strong uterine contractions, facilitating the passage of the fetus through the cervix and vaginal canal^11,35–37^. Immediately following delivery, the uterus undergoes a process of involution, facilitated by active contractions and passive elastic recoil, to prevent maternal hemorrhage and allow the uterus to return back to its nonpregnant state^13,38,39^. Remarkably, the mechanical function of the perimetrium and how it changes in pregnancy is presently unknown.

The uterus performs both active and passive biomechanical functions throughout the menstrual cycle and in pregnancy, shaped in part by tissue architecture and cell composition^13,19,20^. Not only can cells contribute to the active biomechanics of a tissue via contractility, but they can also respond to mechanical stimuli in their surrounding microenvironment through the process of mechanotransduction^40,41^. Elucidating the passive material properties of the tissue is a necessary first step for comprehensive understanding of uterine biomechanics and mechanobiology under normal and pathological states^42,43^. Previous research has focused almost exclusively on characterizing the biomechanics of the myometrium tissue layer at multiple length scales and under different loading conditions^44–48^. Far fewer studies have investigated the material properties of the endometrium and none have investigated the properties of the perimetrium and third-trimester decidua in humans^49^. Further, limited research has been done to evaluate the time-dependent material properties of the uterus, including viscoelasticity and permeability, which also hold the potential to modify cell behavior independent of stiffness^48,50–52^. We broadly hypothesize that the material properties of uterine tissue contribute significantly to the proper initiation and progression of pregnancy, and alterations to normal uterine properties may lead to an array of obstetric and gynecologic pathologies. In order to facilitate the development of physiologically accurate models of the human uterus to address these open questions, this exploratory study seeks to establish the fundamental material properties of all three uterine layers in nonpregnant and pregnant states. Further, this study investigates whether the material properties of all uterine layers are altered in cases pathologic pregnancies, specifically placenta accreta.

## 2. Materials and Methods

### 2.1 Sample Collection

In accordance with IRB approval at Columbia University Irving Medical Center (CUIMC) following written informed patient consent, human uterine tissues were collected from nonpregnant (NP) and pregnant (PG) individuals away from sites of known pathology. A summary of patient clinical characteristics is listed in Table 1 with detailed patient information found in Supplementary Table 1. NP individuals (*n = 6*) underwent total hysterectomies for a variety of gynecologic pathologies including uterine fibroids, endometriosis, adenomyosis, and prolapse. Pregnant uterine tissue was collected in the third trimester for patients undergoing (i) term cesarean sections (PG-CS, *n = 7*) and (ii) cesarean hysterectomies for placenta accreta (PG-A, *n = 6*). All NP and PG tissues were collected at the anterior region of the uterus and contained all three uterine layers, with the exception of three PG-CS samples. A subset of NP and PG-A patient samples was collected at the anterior, posterior, and fundus regions. Tissues were immediately flash-frozen on dry ice and stored at –80°C until testing.

**Table 1.** Summary of Patient Information. Data are presented as median (range). †Gravidity = total number of pregnancies including current; ‡CS = cesarean section; §Hyst = hysterectomy.

| Patient Group | No. Patients | Maternal Age (yrs) | Gestational Age (wks.dys) | No. Previous CS<sup>‡</sup> | Gravidity<sup>†</sup> | Surgery |
| --- | --- | --- | --- | --- | --- | --- |
| NP | 6 | 43.5<br>(36 – 47) | N/A | 0<br>(0 – 1) | 2<br>(0 – 4) | Hyst<sup>§</sup> |
| PG,<br>CS | 7 | 37<br>(29 – 43) | 39.1<br>(37 – 39.3) | 1<br>(0 – 4) | 3<br>(1 – 9) | CS<sup>‡</sup> |
| PG,<br>Accreta | 6 | 35.5<br>(28 – 46) | 35.4<br>(32 – 36.3) | 1.5<br>(1 – 3) | 5<br>(4 – 14) | CS Hyst<sup>‡§</sup> |

### 2.2 Histology

Uterine cross-sections containing all three tissue layers were fixed in 10% formalin solution for 24 hours, subsequently transferred to 70% ethanol solution, paraffin-embedded, and sectioned to a thickness of 5 µm. To observe gross tissue morphology and the distribution of collagen and smooth muscle, all samples were stained for Masson’s trichrome by the Molecular Pathology Core Facilities at CUIMC using standard protocols. Samples were imaged under brightfield microscopy at 10x magnification with a Leica Aperio AT2 whole slide scanner and regions of interest were selected with the ImageScope software (Leica Microsystems, Wetzler, Germany).

#### 2.2.1 Image Quantification

The relative proportions of collagen and smooth muscle were quantified for myometrium tissue stained with Masson’s Trichrome. A custom Matlab (Mathworks, Natick, MA, USA) code utilizing a thresholding function was implemented for such quantification (Fig. 1C). All parameters were kept constant across patient groups. Approximately three to five representative regions of myometrium were used for analysis. Regions containing blood vessels in more than fifty percent of the image area were avoided. To determine the thickness of the perimetrium, measurements were done manually with ImageJ (NIH, Bethesda, MD, USA) on imaged tissue sections stained with Masson’s Trichrome (Fig. 1D).

### 2.3 Nanoindentation

Spherical nanoindentation (Piuma, Optics11Life, Amsterdam, NE) was utilized to determine the time-dependent material properties of uterine tissue (Fig. 2A). A 50 µm probe radius with a cantilever stiffness of 0.15 – 0.5 N/m was used. In preparation for testing, samples were dissected into distinct uterine layers, adhered to a glass dish with superglue (Krazy Glue, Atlanta, GA), and swelled at 4°C overnight in 1X PBS solution. Immediately prior to testing, the sample was equilibrated to room temperature for 30 minutes and subsequently tested in Opti-free contact lens solution (Alcon, Fort Worth, TX, USA) to reduce adhesion between the glass probe and sample^53^. Tissues were indented to a fixed depth of 4 µm under displacement control, corresponding to a 5% indentation strain and contact area of 380 µm^54^. Following a 2 s ramp to the prescribed indentation depth, the probe’s position was held for 15 s to yield a load relaxation curve approaching equilibrium. To ensure reliable measurements taken for thinner tissue layers, the surface of the endometrium, decidua, and perimetrium were directly tested (Fig. 1A). The myometrium was tested orthogonal to the uterine cavity surface in the center of the tissue. Given the irregular size and geometry of the tissues, the number of indentation points varied. At least 100 points were measured to capture intra-sample variability. The distance between individual indentations was kept constant at 200 µm. All tissue sections were at least 1 mm thick and tested within two freeze-thaw cycles.

**Fig. 2.**
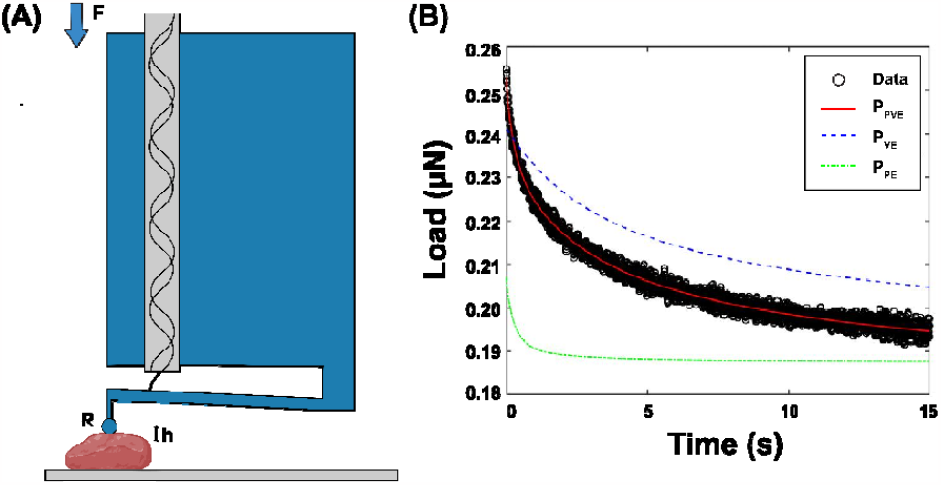
Experimental Approach. **(A)** Diagram of nanoindentation testing (Adapted from Optics11 Life). A spherical probe with radius (R) is attached at the end of a cantilever and indented into the sample at a fixed depth (h), recording load (F) over time. **(B)** Representative load relaxation plot generated from nanoindentation testing in displacement control and fitted with the combined poroelastic-viscoelastic (PVE) material model (solid red line = combined PVE model fit, dotted blue line = viscoelastic material response, solid green line = poroelastic material response).

### 2.4 Data Analysis

In order to determine the time-dependent material properties of the human uterus, load relaxation curves were fit with an established combined poroelastic-viscoelastic (PVE) model in Matlab with a nonlinear least-squares solver (Fig 2B)^54–56^. The following material parameters were determined from the PVE model: (i) stiffness (*E*), (ii) viscoelastic ratio (*E*_∞_/*E*_0_), (iii) intrinsic permeability (*k*), and (iv) diffusivity (*D*).

Fitted data points were excluded from the final data set if the load relaxation curve displayed (i) sharp discontinuities, (ii) increasing loads over time, (iii) Δ*P* (*P*_max_ – *P*_min_) less than 0.005.

The coupled effect of the material’s poroelastic (*P*_*PE*_) and viscoelastic (*P*_*VE*_) force responses is defined as:

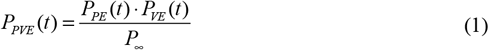

where P_∞_ is the equilibrium force. The viscoelastic force response is calculated using a generalized Maxwell model, consisting of a linear spring connected in parallel with n number of Maxwell units, containing a linear spring and dashpot connected in series. The viscoelastic component of the model is defined by the following equation:

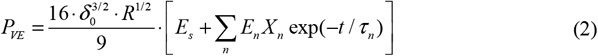

where *E*_*s*_ and *E*_*n*_ are the elastic moduli of the linear spring and the n^th^ Maxwell element (*n = 2*), respectively, δ_*0*_ is the applied indentation depth, *R* is the probe radius, and τ_*n*_ is the characteristic relaxation time of the n^th^ Maxwell element (*n = 2*). A ramp correction factor (*X*_*n*_ = (*τ* _*n*_ / *t*_*r*_) ·[exp(−*t*_*r*_ / (*τ* _*n*_ −1)]) is included to account for the 2 s ramp time (*t*_*r*_) since the original Maxwell model assumes a step loading function. The poroelastic force response is calculated from the analytical solution published in Hu et al. 2010:

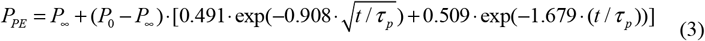

*P*_*0*_ is the initial force at the beginning of the load relaxation curve and is calculated with the Hertzian contact model 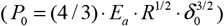 where *E*_*a*_ is the apparent elastic modulus. *P*_∞_ is the equilibrium force given by *P*_∞_ = *P*_0_ / [2(1−*ν* _*d*_)] where ν_*d*_ is the drained Poisson’s ratio. τ_*p*_ is a fitted constant in the model defined as:

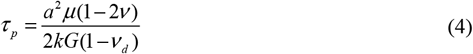

where *a* is the contact radius 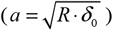, μ is the interstitial fluid viscosity (μ = 0.89 × 10^−3^ Pa·s), ν is Poisson’s ratio (ν = 0.5), *k* is intrinsic permeability, and *G* is the shear modulus. Elastic modulus (*E*) is calculated from shear modulus as *E* = 3*G*. Diffusivity (*D*) is calculated as *D* = *a*^2^ / *τ* _*p*._

### 2.5 Statistical Analysis

For image quantification data, statistical analysis was performed in Prism 9.4.0 (GraphPad, California, USA). Multiple nested t-tests were utilized to analyze collagen/smooth muscle content and perimetrium thickness. Due to the complexity of the dataset, analysis of material parameters was conducted in RStudio version 1.3.1056. The normality of the data was first assessed with a QQ-plot. Stiffness, permeability, and diffusivity data were normalized with a logarithmic transformation. No transformation was made for viscoelastic ratio. All parameters were subsequently analyzed with a linear mixed model, assessed by (1) pregnancy state accounting for patient ID and (2) tissue group accounting for patient ID. Statistical significance was set at a 95% confidence level for all analyses. P-value symbols follow a standard GraphPad (GP) style (ns: p > 0.05; ^*^ p ≤ 0.05 ; ^**^ p < 0.01 ; ^***^ p < 0.001; ^****^ p < 0.0001) with an added distinction for trends (p ≤ 0.1) denoted by a # symbol.

## 3 Results

### 3.1 Distinct Layers of the Human Uterus

Histological images of the human uterus demonstrate clear morphological and compositional differences across all tissue layers in NP and PG states (Fig. 1B). The NP endometrium of all patient samples contains glandular epithelium and stromal cells embedded in a collagen-dense extracellular (ECM) matrix (Fig. 1B). The approximate menstrual cycle stage for each of the NP individuals is noted in Table S2. PG decidua tissue largely contains decidualized stromal cells in a collagenous matrix with thin or non-visible glandular epithelium. No notable differences are present between the decidua of PG-CS and PG-A samples, which represent the decidua parietalis, a region of tissue away from the site of placentation (Fig. 1B). Within a mm^2^ region of tissue, the PG myometrium exhibits a significant increase in the size of smooth muscle fibers and a relative decrease in collagen content when compared to the NP myometrium (Fig. 1C). No discernible changes in tissue morphology or composition are evident between the PG-CS and PG-A samples (Fig. 1B). Lastly, the perimetrium is observed to be a thin layer of tissue composed primarily of collagen that appears to thicken in pregnancy (Fig. 1D). Greater variability in the perimetrium thickness of PG-A samples is noted (Fig. 1D).

### 3.3 Stiffness of the Human Uterus

Overall, uterine stiffness (*E*) ranges from 10^2^ – 10^3^ Pa, with a large degree of spatial heterogeneity observed for all tissues measured (Fig. 3C, S6). Notable stiffness differences are evident across uterine layers and as a result of pregnancy (Fig. 3). For the NP uterus, the myometrium is significantly and consistently stiffer than the endometrium for all patients (Fig. 3A, S5) and the perimetrium in four out of six evaluated (Fig. 3A, S5). Overall, the NP perimetrium exhibits a decreasing trend (*p = 0*.*096*) in stiffness relative to the NP myometrium (Fig. 3A). In pregnancy, the PG-CS myometrium is significantly stiffer than the PG-CS decidua. However, for PG-A samples, this stiffness difference is only a trend *(p = 0*.*073)* (Fig. 3A, S5). For both PG-CS and PG-A tissue samples, there is no significant difference in tissue stiffness between the myometrium and perimetrium (Fig. 3A, S5).

**Fig. 3.**
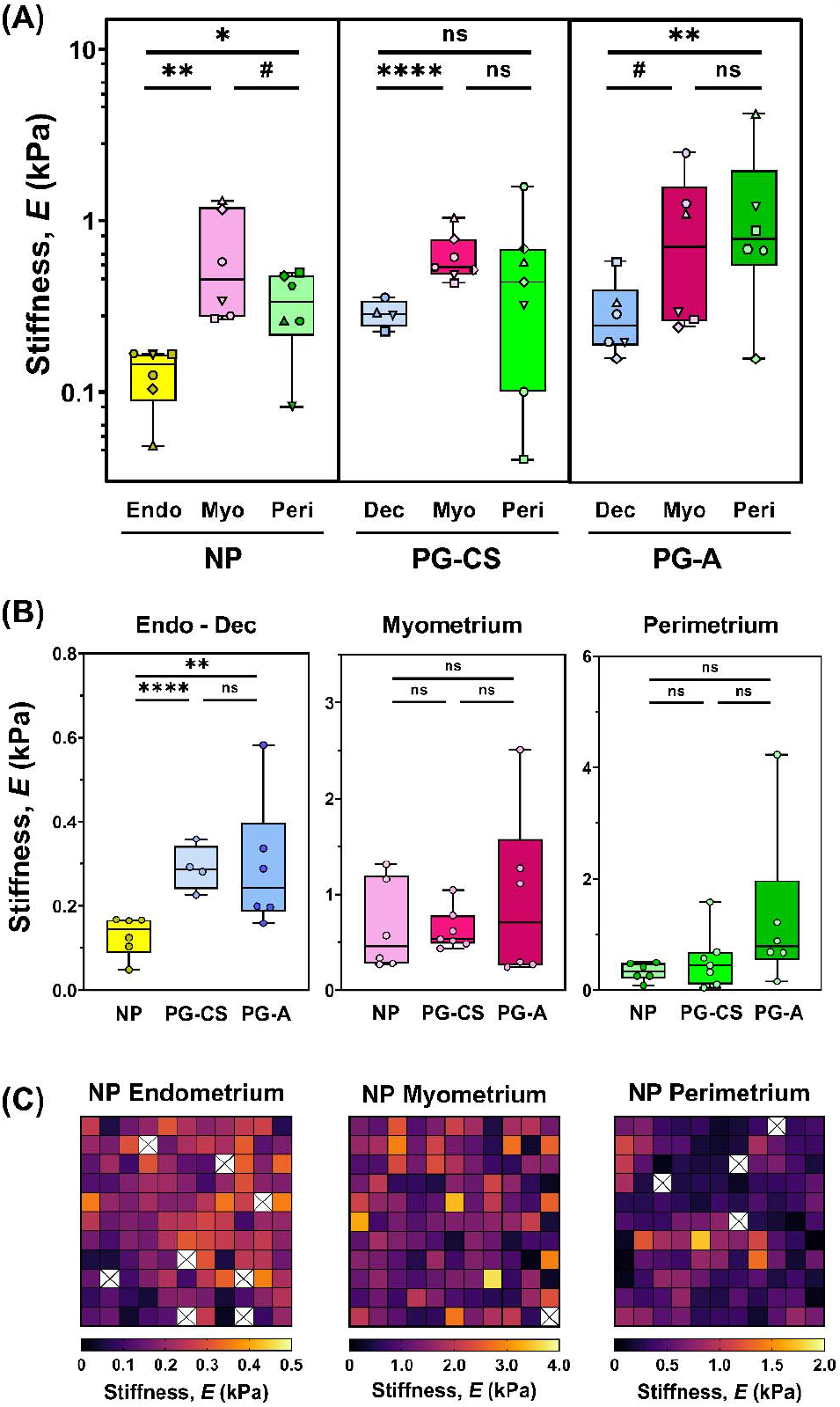
Stiffness of Human Uterine Layers. **(A)** Summary of stiffness values for each tissue layer at the anterior region organized by patient groups: nonpregnant (NP), cesarean section pregnancy (PG-CS), and placenta accreta pregnancy (PG-A). Each distinct symbol represents the median value of all indentation points measured for a single patient. Data are presented as box and whisker plots on a log_10_ scale. **(B)** Stiffness values separated by tissue layer, compared across all three patient groups, plotted on a linear scale. Statistical significance is denoted by the following symbols: ns, p > 0.05; # p ≤ 0.1; ^*^ p ≤ 0.05 ; ^**^ p ≤ 0.01; ^****^ p ≤ 0.0001). **(C)** Representative stiffness heatmaps for NP uterine layers. Grid of 11 × 11 indentation points corresponds to a 2 mm x 2 mm region. [X] indicates points removed due to exclusion criteria.

Comparing across all three patient groups, the PG decidua is stiffer than the NP endometrium in third trimester (Fig. 3B). Greater variability in decidua stiffness is noted for PG-A tissue samples (Fig. 3, S5). No change in the stiffness of the myometrium is observed between the NP and PG tissue samples (Fig. 3B). Additionally, there is no statistically significant difference in perimetrium stiffness between the NP and PG tissue samples, yet the PG-A perimetrium layer exhibited the stiffest points measured for the human uterus (Fig. 3B). It is important to note that these extrema in PG-A stiffness values measured for all three tissue layers cannot be considered true outliers as they represent measurements from two distinct patients away from any visual pathology (Fig. 3B).

### 3.4 Regional Variations in Uterine Layer Stiffness

To investigate whether uterine stiffness varies across anatomic regions, full-thickness sections of the uterus were collected from a subset of NP and PG-A patients at the anterior, posterior, and fundus regions. On an individual patient basis, variations in stiffness exist for all NP and PG uterine layers (endometrium-decidua, myometrium, and perimetrium) across the three anatomic regions investigated (Fig. S1, S2). Given the relatively few number of patients included in this analysis, it is unclear whether systematic variations exist for any particular anatomic region (Fig. S1). Interestingly, the proximity of the PG-A uterine layers to the site of placentation, as noted in Table S1, does not correlate with any regional increases in stiffness in this dataset.

### 3.5 Time Dependent Material Properties of the Human Uterus

Viscoelastic ratio (*E*_∞_/*E*_0_) is defined as the ratio between the equilibrium and instantaneous elastic moduli and reflects a material’s degree of viscoelasticity (0 = viscous fluid, 1 = elastic solid). Median viscoelastic ratio values of all uterine layers across NP and PG tissue samples are between 0.4 and 0.6, indicating that the human uterus behaves as both a liquid and a solid at the microscale (Fig. 4). Interestingly, a difference in the viscoelastic ratio is only observed for the endometrium-decidua layer, which forms the basis of the maternal-fetal interface (Fig. 4). Relative to the NP endometrium, the viscoelastic ratio of PG-CS and PG-A decidua increases (Fig. 4B). Therefore, this finding suggests that the endometrium-decidua becomes less viscous in pregnancy. Both the NP endometrium and PG-CS decidua exhibit smaller viscoelastic ratios relative to both the myometrium and perimetrium, however, this trend is not observed for the PG-A tissue samples (Fig. 4A).

**Fig. 4.**
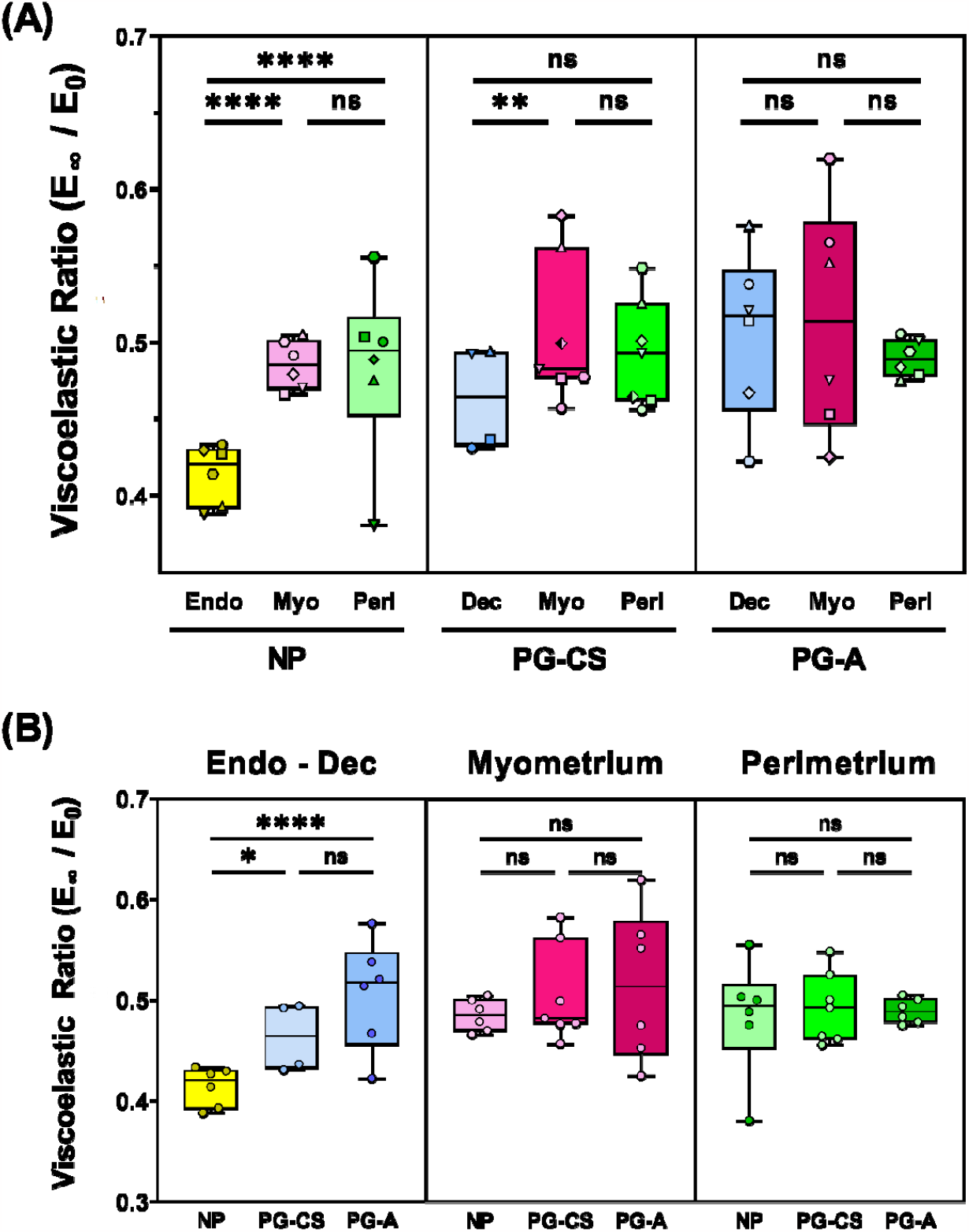
Viscoelasticity of Human Uterine Layers. Summary of viscoelastic ratio values for each tissue layer at the anterior region organized by patient group (NP, PG-CS, PG-A). Each distinct symbol represents the median value of all indentation points measured for a single patient. Data are presented as box and whisker plots on a log_10_ scale. **(B)** Viscoelastic ratio values separated by tissue layer, compared across all three patient groups, plotted on a linear scale. Statistical significance is denoted by the following symbols: ns, p > 0.05; ^*^ p ≤ 0.05 ; ^**^ p ≤ 0.01; ^****^ p ≤ 0.0001).

Intrinsic permeability (*k*) describes the motion of molecules through a material due to physical pore geometry and is a fitted parameter from the poroelastic component of the PVE material model. Median permeability values range from 10^−17^ to 10^−15^ m^2^ for all uterine layers in both NP and PG tissue samples (Fig. 5). The NP endometrium exhibits decreased permeability relative to the NP myometrium and perimetrium. Yet, no differences are observed between NP and PG tissue samples (Fig. 5, S4). Further, no change in permeability exists across the PG-CS and PG-A uterine layers (Fig. S4). Lastly, by taking the square root of the permeability values, we determine that the mean pore size for all uterine layers is on the order of 10 to 100 nm.

**Fig. 5.**
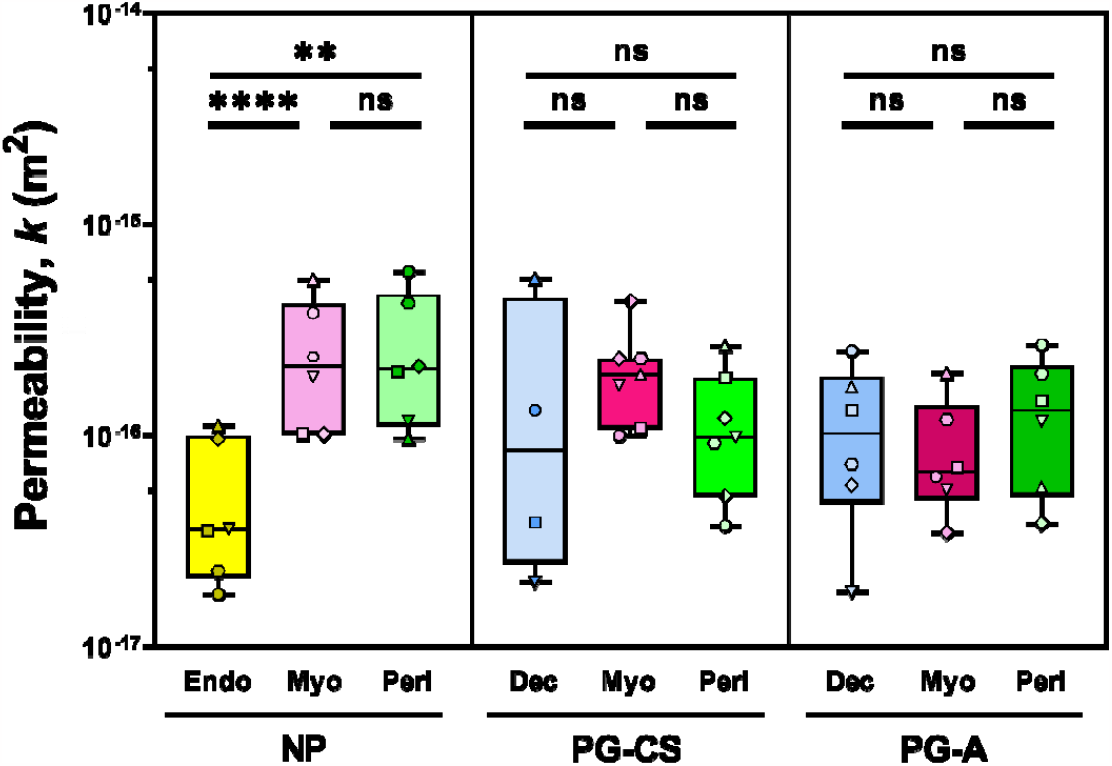
Permeability of Human Uterine Layers. Summary of permeability values across uterine layers at the anterior region for each patient group (NP, PG-CS, PG-A). Each distinct symbol represents a single patient for which tens to hundreds of indentation points have been averaged. Data are presented as box and whisker plots on a log_10_ scale. Statistical significance is denoted by the following symbols: ns, p > 0.05; ^**^ p ≤ 0.01; ^****^ p ≤ 0.0001).

The final time-dependent material property determined from the PVE model is diffusivity (*D*). This parameter describes the motion of molecules through a material over time and is calculated from values of permeability and stiffness. Diffusivity exhibits similar trends to permeability and viscoelastic ratio, with median values ranging from 10^−12^ to 10^−8^ to m^2^/s for all uterine tissues (Fig. S3). Diffusivity increases in pregnancy for the PG decidua relative to the NP endometrium, with no changes observed for the myometrium and perimetrium in pregnancy (Fig. S3B). Both the NP endometrium and PG-CS decidua exhibit smaller diffusivity values relative to patient-matched myometrium, yet this difference is not present for PG-A tissue samples (Fig. S3A). It is important to note that because this parameter is calculated, relative changes in diffusivity across tissue layers and pregnancy state are dependent on stiffness and permeability values. A summary of all material properties can be found in Table 2.

**Table 2.**
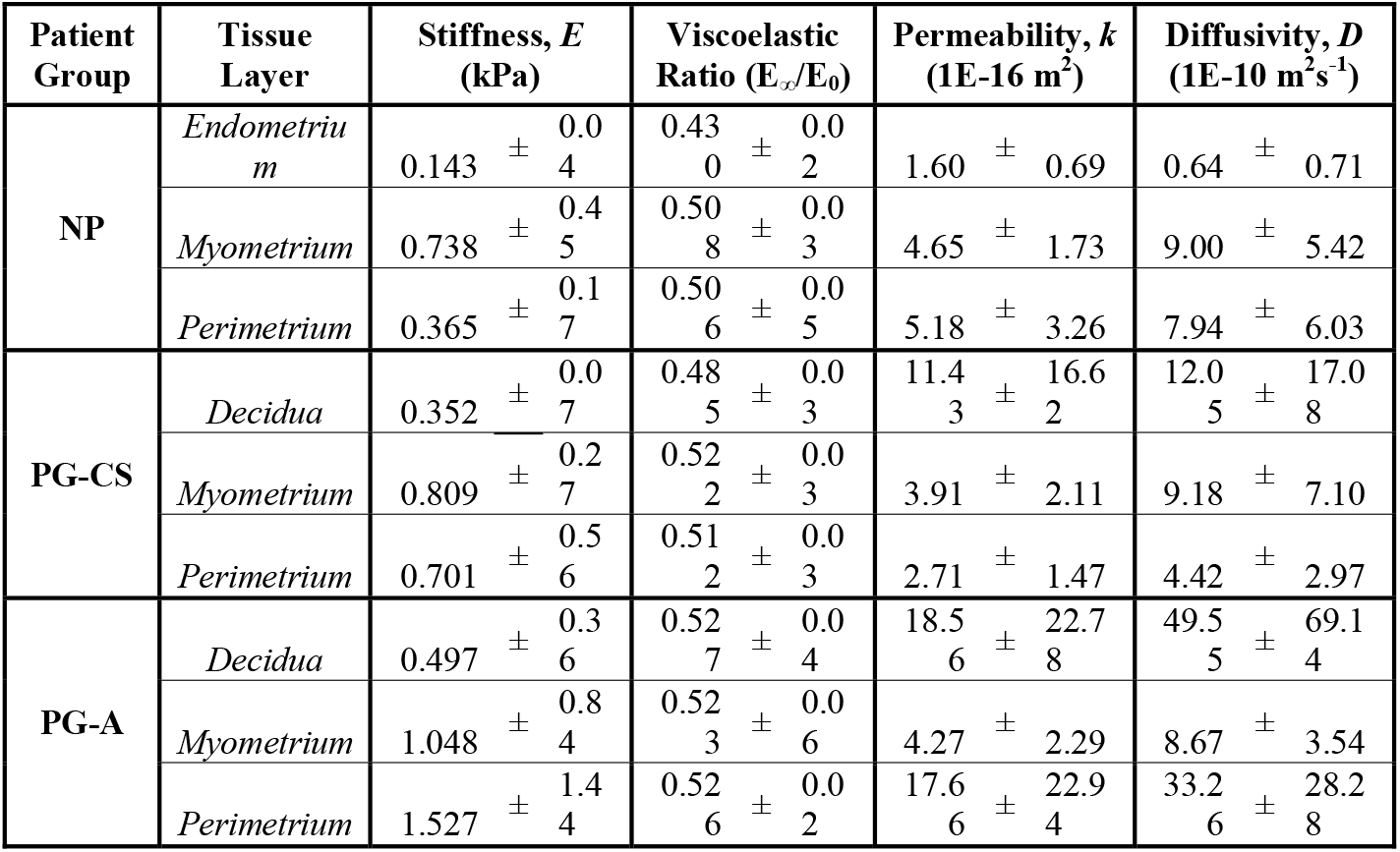
Summary of Material Properties of Human Uterine Layers. Mean ± std. dev calculated for each tissue layer across all samples for a given patient group.

## 4. Discussion

This study represents a first-of-its-kind observational study characterizing the material properties of human uterine tissue layers and the changes, or lack thereof, that result from pregnancy. The first measurements of uterine permeability and diffusivity for all tissue layers are reported in this study and include the first stiffness measurements of the perimetrium and third-trimester PG decidua. Such data are critical for the development of physiologically accurate *in vitro* and *in silico* models of the uterus in the context of pregnancy and gynecological disorders^57^. A probe radius of 50 µm (100 µm diameter) was specifically chosen to match the size and contact area of the invading embryo^11^, providing measurements that are directly relevant for mechanobiology studies on embryo implantation and placentation.

### 4.1 Material Behavior of Human Uterine Layers

Fundamental differences in the tissue composition and structure of all three uterine layers are reflected in the measured material properties. Changes in stiffness and time-dependent material properties of the uterus are observed in both NP and PG tissue samples, with the most remarkable differences seen in the endometriumdecidua layer. In nonpregnancy, the endometrium exhibits decreased stiffness, viscoelastic ratio, permeability and diffusivity relative to both the myometrium and perimetrium (Fig. 3A, 4A, S3A). Mucous production by luminal and glandular epithelial cells in the endometrium is one possible explanation for the existence of a tissue that is more viscous and less permeable^11,58,59^. In pregnancy, the decidua is less stiff than the myometrium, however, differences relative to the perimetrium are variable (Fig, 3A). The degree to which this is influenced by confounding variables such as gynecological disorders, age, gravidity, and the site of placentation is unknown.

### 4.2 Effect of Pregnancy on Uterine Remodeling

A clear increase in the stiffness, viscoelastic ratio, and diffusivity of the endometrium-decidua layer is observed in pregnancy (Fig. 3B, 4B, S3B). This change in material properties indicates that the maternal-fetal interface undergoes significant remodeling in pregnancy, which is confirmed by histological findings (Fig. 1B). The process of endometrial decidualization leads to glandular epithelium hypotrophy and stromal cell differentiation which in turn contribute to ECM-level changes^60^. The endometrium-decidua is largely made up of collagen, exhibiting notable shifts in the amount, size and subtypes present in pregnancy^14,61–63^. A bulk of the relevant research has been conducted on animal and first-trimester human tissues with comparatively less known for the third-trimester human decidua. Mechanically, a striking paucity of data has been published on the material properties of the human endometrium and decidua. A key study by Abbas et al. (2019) reported the first stiffness measurements of these tissues, demonstrating no difference in stiffness between the nonpregnant endometrium and decidua parietalis of first-trimester pregnancies^49^. The data presented in our study illustrates clear stiffening of the third-trimester decidua parietalis relative to NP endometrium in both pregnant patient cohorts (Fig. 3B). This key finding highlights the significant remodeling the maternal-fetal interface undergoes from the first to third trimesters. It is presently unknown whether this shift in material properties results from biological factors, mechanical triggers, or the interplay of both. It is well established that dramatic fetal growth is observed from the first trimester until parturition, with a majority occurring in the third trimester (*33-36*). We, therefore, hypothesize that increased mechanical loads on the uterus throughout pregnancy are responsible for decidual stiffening, thereby suggesting that this tissue is mechanoresponsive.

Dramatic growth and remodeling of the pregnant myometrium are necessary to accommodate fetal growth and prepare the uterus for coordinated labor contractions^11,13,34^. Interestingly, this study showed no significant difference in the material properties of the gravid myometrium relative to nonpregnancy when measured with nanoindentation (Fig. 3B, 4B, S3B, S4). This finding is intriguing in the context of observed histological changes to the pregnant myometrium. Notably, we find a decrease in the relative proportion of collagen to smooth muscle content in the pregnant myometrium (Fig. 1C). Yet, given the immense amount of growth the myometrium undergoes in gestation, the total amount of collagen has been previously shown to increase in term pregnancies ^61^. Elucidating how the structure and composition of the uterus changes across length scales and are reflected in material property measurements are critical for understanding pregnancy in its totality. On the microscale, the material properties measured in this study by nanoindentation are vital for understanding fundamental cell-ECM interactions. Within the large-strain regime, macroscale material property changes, captured with tension and compression, provide a broader understanding of how ECM fiber bundles, smooth muscle cell fascicles, and the supporting ground substance contribute to uterine remodeling. Parallel work by our group has recently evaluated the macroscale material properties of the myometrium in an overlapping patient cohort^65^. Under small strains, no change in myometrium stiffness was found in pregnancy with indentation and tension tests, showing agreement with the nanoindentation data presented in this paper^65^. Material changes to the PG myometrium were only observed for strains above 30%, demonstrating increased tissue extensibility relative to the NP uterus^65^. Collectively, these data highlight the functional role of the uterus in pregnancy, where the uterus must remain structurally intact despite continuous growth and remodeling to accommodate the increasing weight and size of the fetus.

Lastly, this study shows for the first time that the perimetrium also undergoes considerable remodeling in pregnancy as evidenced by an increased thickness of this collagen-dense layer (Fig. 1D). These compositional changes, however, are not reflected by material property changes to the perimetrium (Fig. 3B, 4B, S3B, S4). Key limitations of the dataset include the small sample size (n = 6) and inherent uterine pathology of nonpregnant and pregnant patients (Table S1). A prospective study powered to assess this difference is needed. Aside from its existence as the outermost layer of the uterus, little is understood regarding the structure and function of this tissue layer, particularly during pregnancy. We hypothesize that the perimetrium provides structural support to the uterus as the primary boundary between the myometrium and the abdominal cavity. Further, we posit that remodeling of the perimetrium is important in the proper development of pregnancy, and without this, obstetric complications such as uterine rupture may arise.

### 4.3 Effect of Placenta Accreta on Uterine Material Properties

No significant differences in the material properties of all uterine layers are observed between the PG-CS and PG-A tissue samples away from the site of placentation and any visible pathology (Fig. 3B, 4B, S3B, S4). Notably, the stiffest measurements in this dataset were taken from PG-A tissue samples (Fig. 3, S6). Given the limited size of this patient cohort, it is unclear whether the material properties of the uterus were altered as a result of invasive placentation or were already transformed by a previous cesarean delivery or the presence of underlying gynecologic pathology. Notably, cesarean section surgeries are known to result in uterine scar tissue, contributing to thin or absent decidua at the incision site, and may predispose an individual to placenta accreta in subsequent pregnancies^13,66^. Differences in the material and structural properties of cesarean scar tissue compared to healthy decidua is one hypothesis for altered trophoblast invasion emblematic of placenta accreta.

Variations in gestational age may also influence the material properties PG samples. All cesearean hysterectomies for the collection of PG-A tissues occurred pre-term, before 37 weeks of gestation, while PG-CS tissue were collected between 37 and 40 weeks of pregnancy. The role of mechanics in the pathophysiology of placenta accreta remains an open question and warrants statistically-powered follow-up study to tease out the effect of disease and patient confounding factors.

### 4.4 Regional Variations in Uterine Stiffness

Regional differences in stiffness for all uterine layers were observed for a subset of NP and PG-A patients (Fig. S1). Previous work by Fang et al. (2021) also suggests regional variations in the mechanical properties of the NP and PG myometrium representing a single patient per group^48^. Collectively, these data represent a small number of patients, making it difficult to determine whether systemic regional changes in uterine stiffness exist. Obstetric and gynecologic history, such as the number of previous pregnancies (i.e., gravidity), previous sites of placentation, number of cesarean sections, and presence of gynecologic pathology, may contribute to regional differences observed across different patients. Limited by the small sample size of this dataset, no correlations between regional stiffness and these confounding variables can be made.

Previous work by Abbas et al. (2019) found increased stiffness of decidua basalis tissue (i.e., at the site of placentation) relative to the decidua parietalis (i.e., away from the placenta) in the first trimester^49^. This result suggests that extravillous trophoblast invasion, which serves as the basis for placenta formation and attachment to the uterus, is responsible for this local stiffening phenomenon ^49^. Interestingly, data generated from our study suggests that the proximity of decidua parietalis tissue to the site of placentation does not correlate with increased stiffness. Future studies are needed to assess stiffness changes in decidua basalis tissue in humans.

### 4.5 Limitations

Patient variability is a critical consideration likely contributing to the wide range of material property values reported in this study. Confounding factors such as age, gravidity, parity, number of previous cesarean sections, region of placentation, and any underlying gynecological conditions may have a significant yet unknown effect. Notably, all nonpregnant tissues measured in this study are inherently pathological, diagnosed with uterine fibroids, prolapse, adenomyosis, and/or endometriosis (Table S1). It is unclear the degree to which, if any, these gynecological disorders impact the composition, structure, and mechanics of the uterus. Although changes in the proportion of collagen and smooth muscle content are observed for the nonpregnant and pregnant myometrium analyzed in this study, additional research is needed to confirm whether this is a normal byproduct of pregnancy or the reflection of a gynecologic pathology in the nonpregnant subjects. It is important to note that all tissues in this study were tested away from any visually identifiable pathology and confirmed with histology to contain all three uterine layers. Moreover, given that the stiffness of the endometrium measured in this dataset matched the values reported by Abbas et al. (2019) for healthy NP endometrium, it is a reasonable assumption that the NP tissue measurements are minimally affected by pathology^49^.

Further, it is important to consider the length scale and technical limitations of nanoindentation testing, which may contribute to unintended biases in the dataset. Given the probe parameters utilized in this study, the contact radius and surface area of each indentation test was approximately 14.14 μm and 355 μm^2^, respectively. On average, eukaryotic cells have diameters ranging from tens to hundreds of microns, with intracellular organelles on the nm-scale^67^. Therefore, the nanoindentation testing employed in our study most likely measures the material properties for a collection of extracellular matrix and cellular components at a given indentation point and is at a scale too large to discern individual intracellular structures. In addition, the overall surface area for each of the three tissue layers evaluated is considerably larger (cm^2^ to m^2^ depending on the stage of pregnancy) than the mm^2^ testing area covered by nanoindentation. To address this limitation and minimize its effect, a large number of indentation points (10^1^ to 10^2^) was chosen to characterize each tissue, totaling more than 7000 individual measurements across all experimental groups.

## 5. Conclusions

The data presented in this study yield key insights into the material properties of the uterus and underscores the mechanical nature of pregnancy. Notable time-dependent material property changes are observed across tissue layers and as a result of pregnancy. Alterations in parameters of stiffness, viscoelastic ratio, permeability, and diffusivity are demonstrated only at the maternal-fetal interface for the endometrium-decidua layer. Overall, the role of mechanics in the initiation and progression of gynecological and obstetric disorders remains an understudied area of research and the data generated from this work will advance *in vitro* and *in silico* models of the uterus and pregnancy.

## Supporting information

Supplemental - Nanoindentation Data

## 7. Acknowledgments

We would like to thank Dr. George Gallos and Dr. Arnold Advincula for their assistance in the collection of uterine tissue and Dr. Qi Yan for statistics consultation. This work is supported financially by the Eunice Kennedy Shriver National Institute of Child Health & Human Development of the National Institutes of Health under award number 1R01HD091153 (KMM), NSF Graduate Research Fellowship (DMF) and the Iris Fund (KMM). The content is solely the responsibility of the authors and does not necessarily represent the official views of the National Institutes of Health.

## 8. Author contributions

Conceptualization: DMF, MLO, KMM

Methodology: DMF, MLO, KMM

Investigation: DMF

Formal analysis: DMF, SRR, JLL, XC

Resources: SF, KMM, JYV

Supervision: MLO, KMM

Writing—original draft: DMF

Writing—review & editing: DMF, SRR, MLO, KMM, JYV, XC

## 10. Data and materials availability

All data are available in the main text or the supplementary materials.

