## Supplemental Figures and Tables for "Material Properties of Nonpregnant and Pregnant Human Uterine Layers"

Daniella Fodera *et al.*

*Corresponding authors.

Michelle Oyen:

Kristin Myers:

**This PDF file includes:**

Figs. S1 to S9

Table S1 to S2

**Other Supplementary Materials for this manuscript include the following:**

Data S1

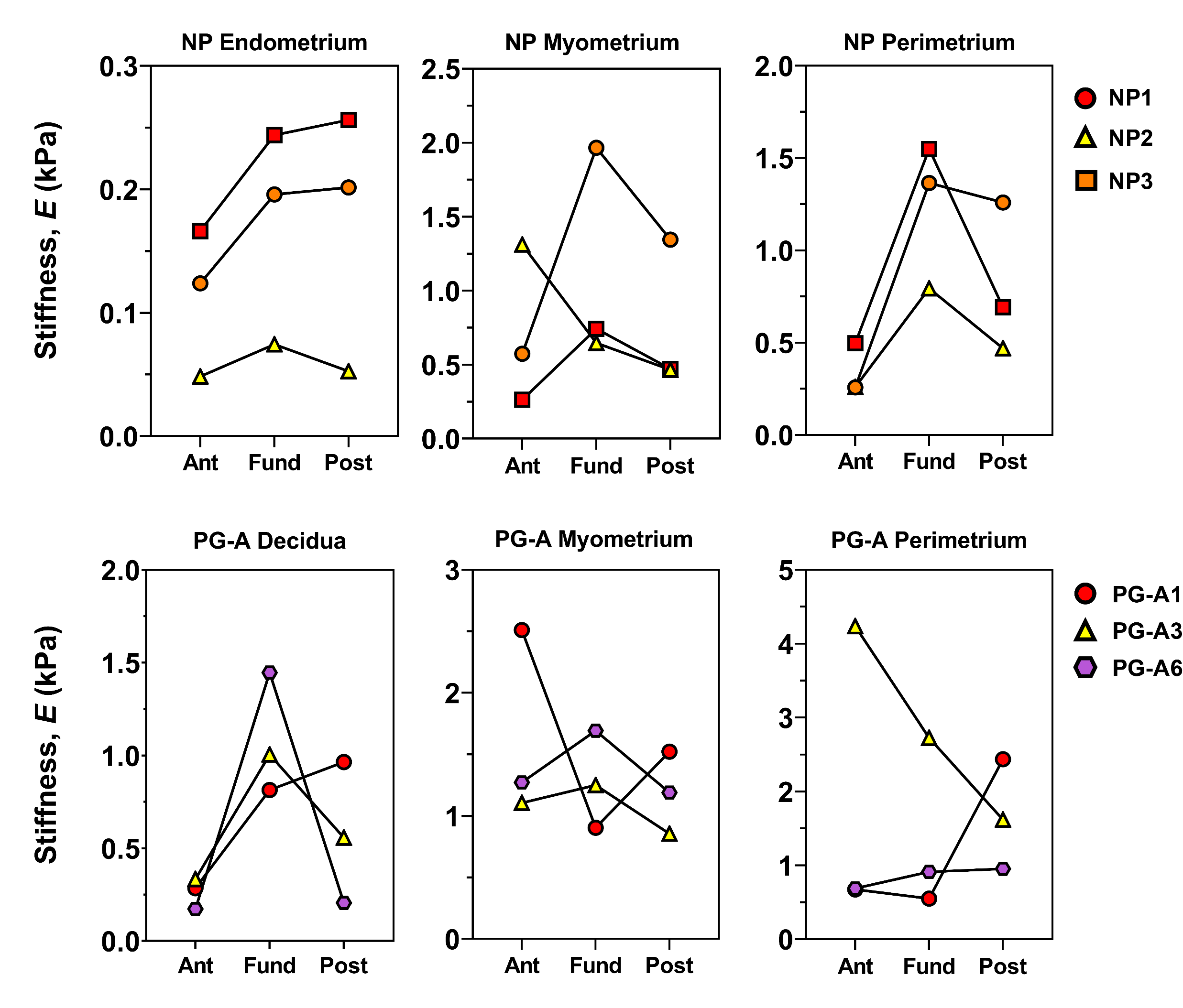

**Figure S1.** **Patient-Matched** **Regional Variations in Uterine Layer Stiffness.** Each connecting line indicates the matched median stiffness values for a single patient across anterior, fundus, and posterior regions. Data are shown for three NP (top row) and PG-A (bottom row) patients and grouped by tissue layer.

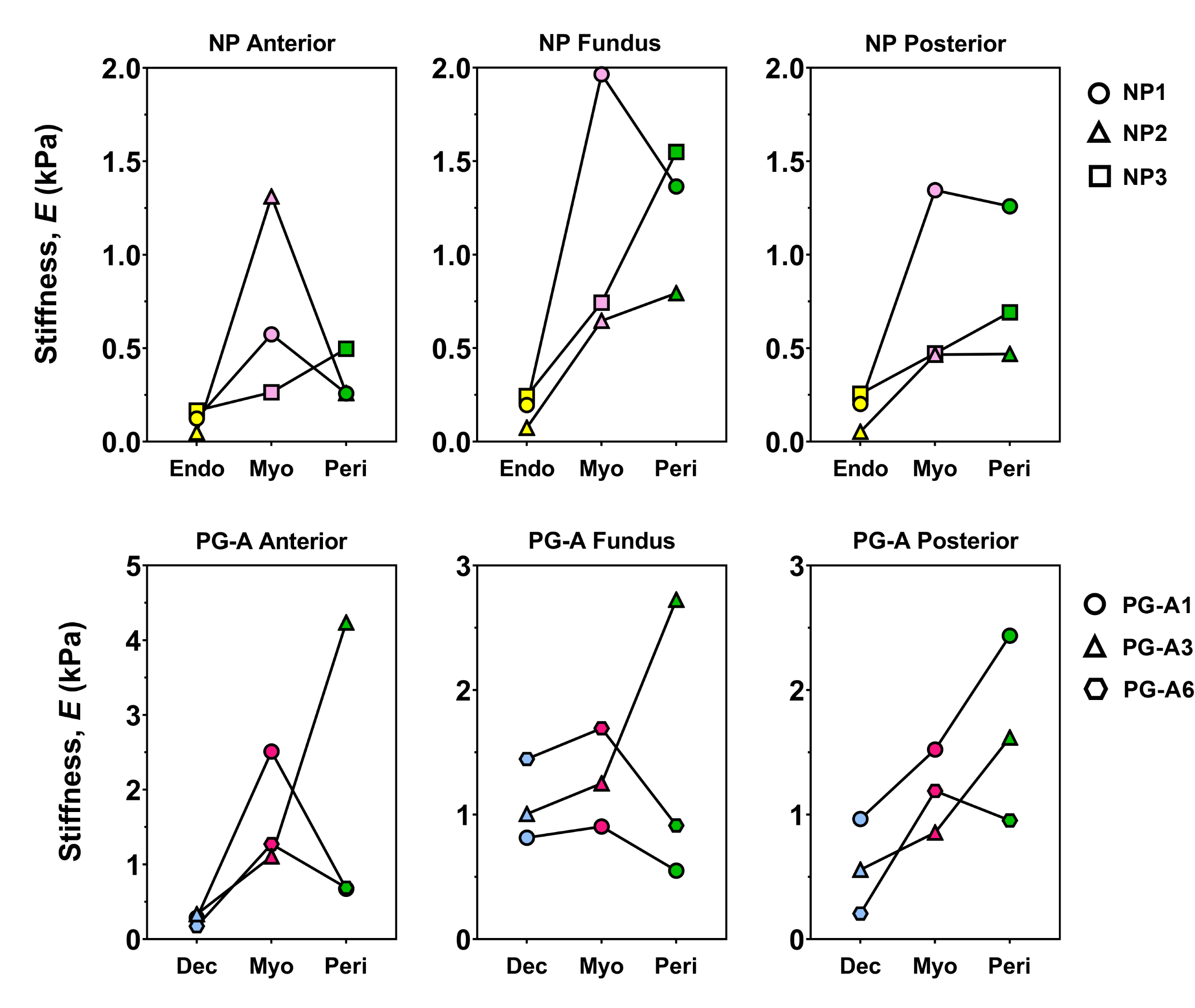

**Figure S2.** **Patient-Matched** **Variations in Uterine Layer Stiffness Grouped by Regions.** Each connecting line indicates the matched median stiffness values for a single patient across anterior, fundus, and posterior regions. Data are shown for three NP (top row) and PG-A (bottom row) patients and separated by tissue layer.

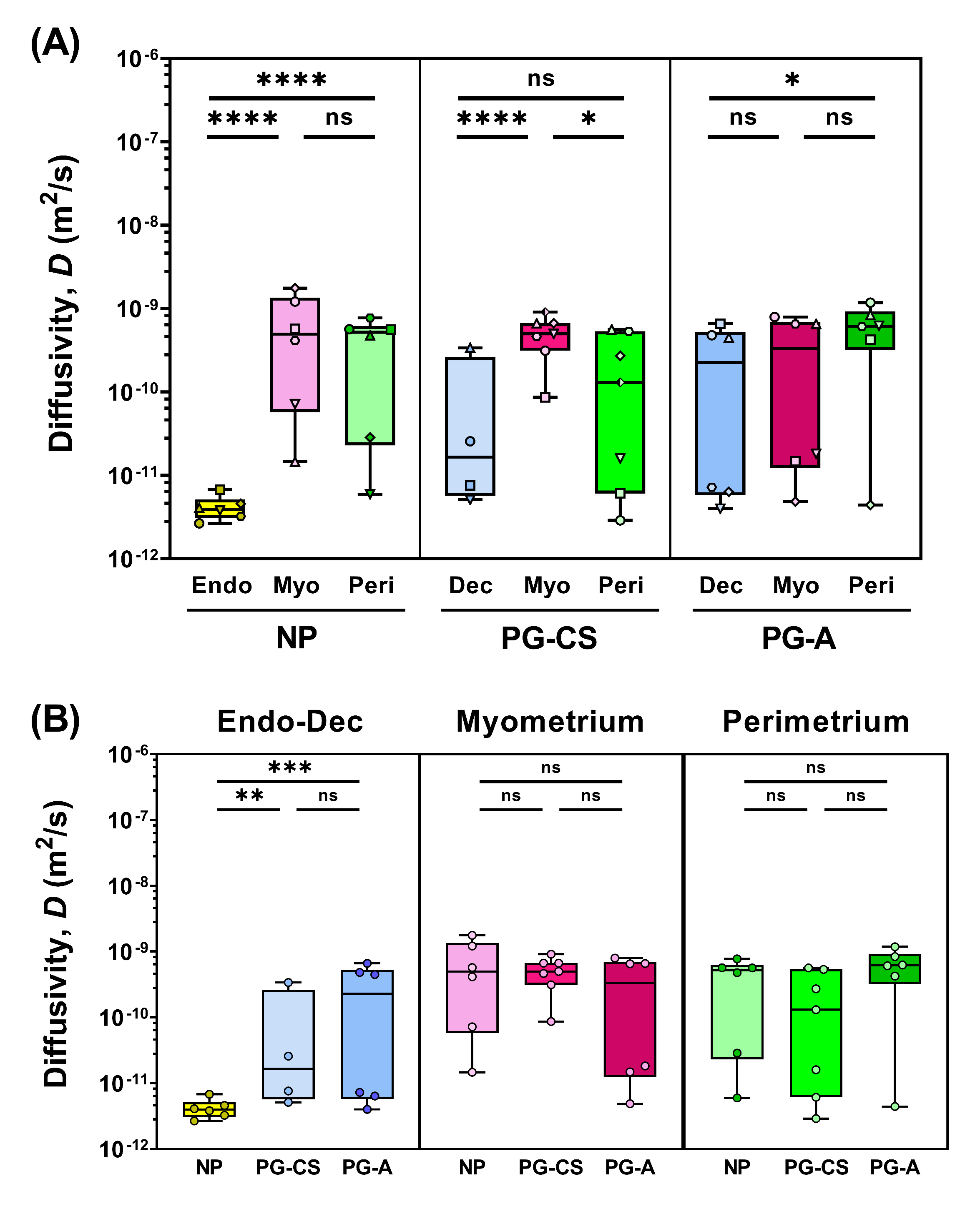

Figure S3. Diffusivity of Human Uterine Layers. (A) Summary of diffusivity values for each tissue layer at the anterior region organized by patient group (NP, PG-CS, PG-A). Each distinct symbol represents the median value of all indentation points measured for a single patient. (B) Stiffness values compared across all three patient groups for a single tissue layer. Data from (A) and (B) are presented as box and whisker plots on a log_10_ scale.

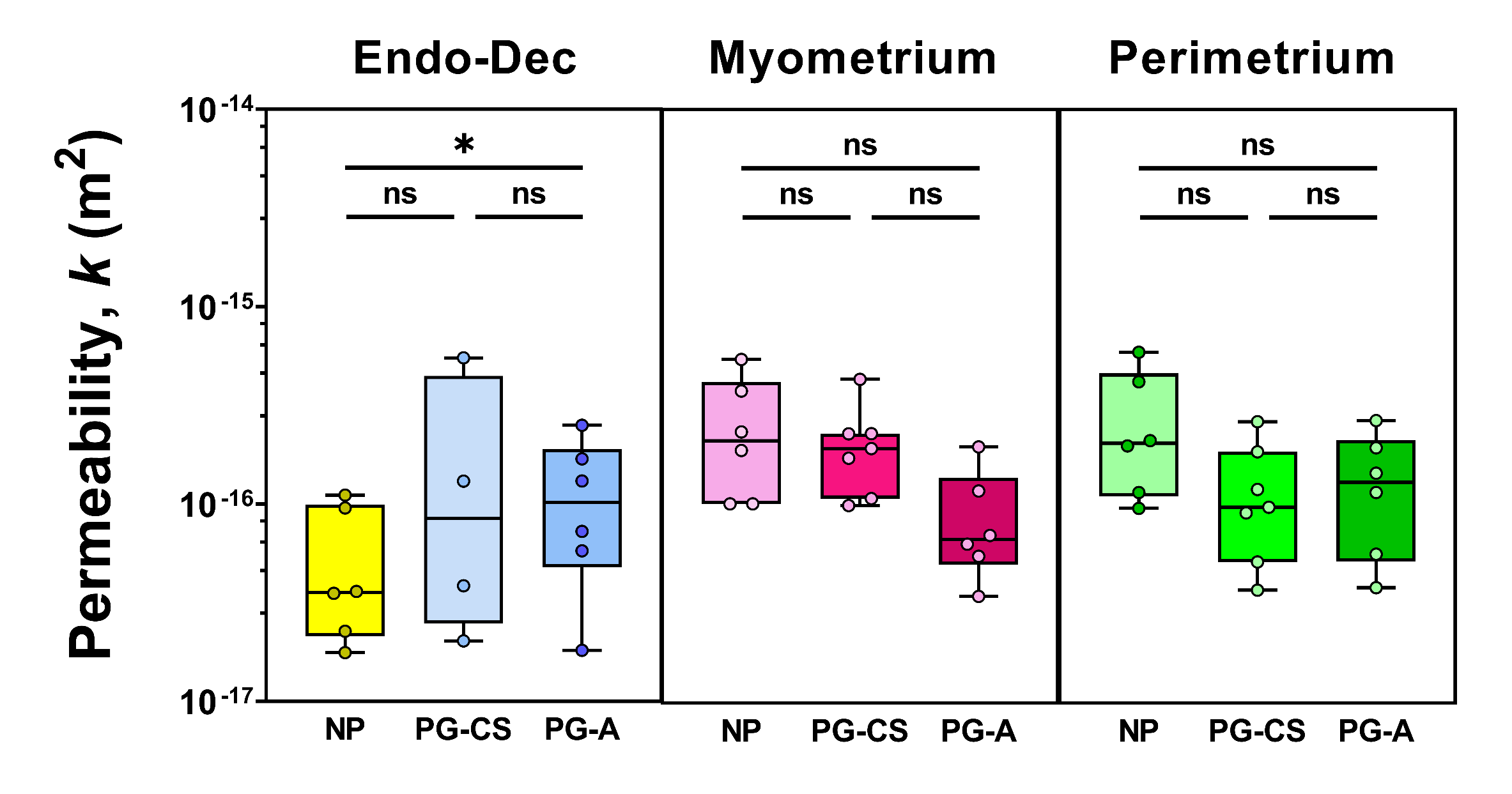

Figure S4. Permeability of Human Uterine Layers Across Patient Groups. Summary of permeability values separated by tissue layer and compared across all three patient groups (NP, PG-CS, PG-A). Each symbol represents the median value of all indentation points measured for a single patient. Data are presented as box and whisker plots on a log_10_ scale.

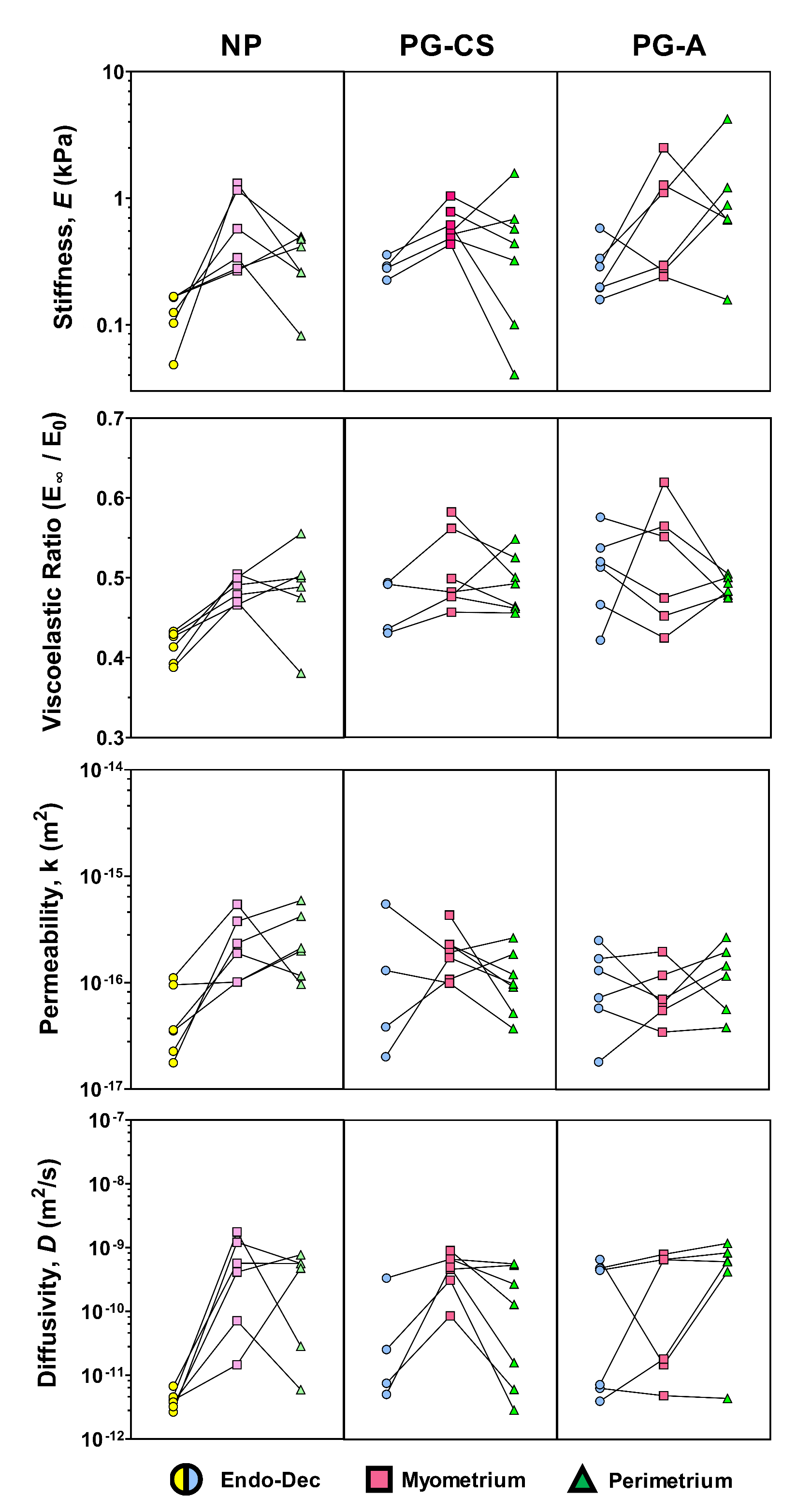

Figure S5. Patient-Matched Material Properties Across Tissue Layers. Each connecting line indicates the matched tissue layers for a single individual for the following parameters: stiffness, viscoelastic ratio, permeability, and diffusivity. Individual symbols represent the median value of all indentation points.

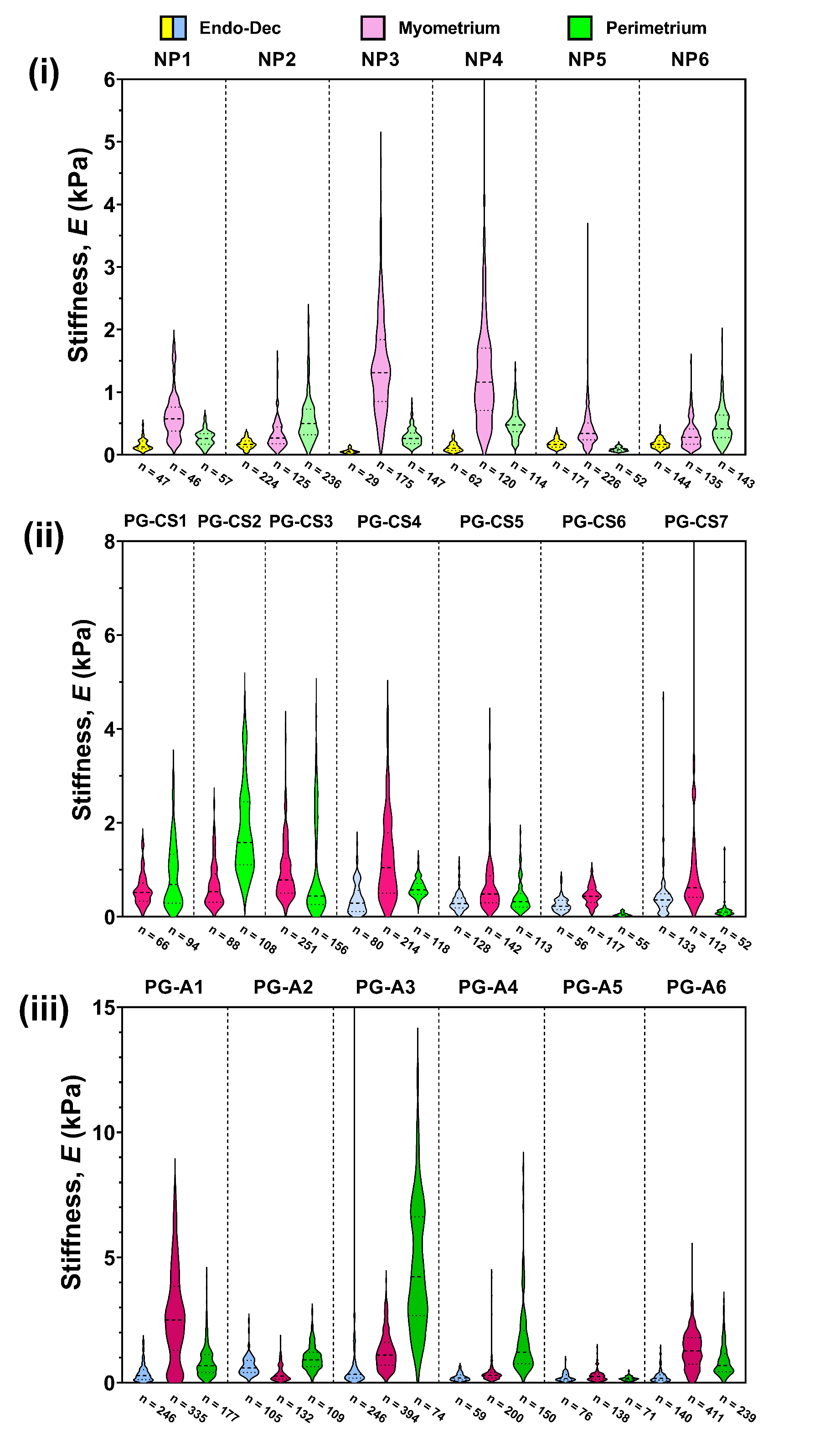

Figure S6. Distribution of Stiffness Values for Individual Patient Uterine Layers. Data are presented as violin plots on a linear scale for each patient group: (i) NP, (ii) PG-CS, (iii) PG-A. ‘n’ denotes technical replicates.

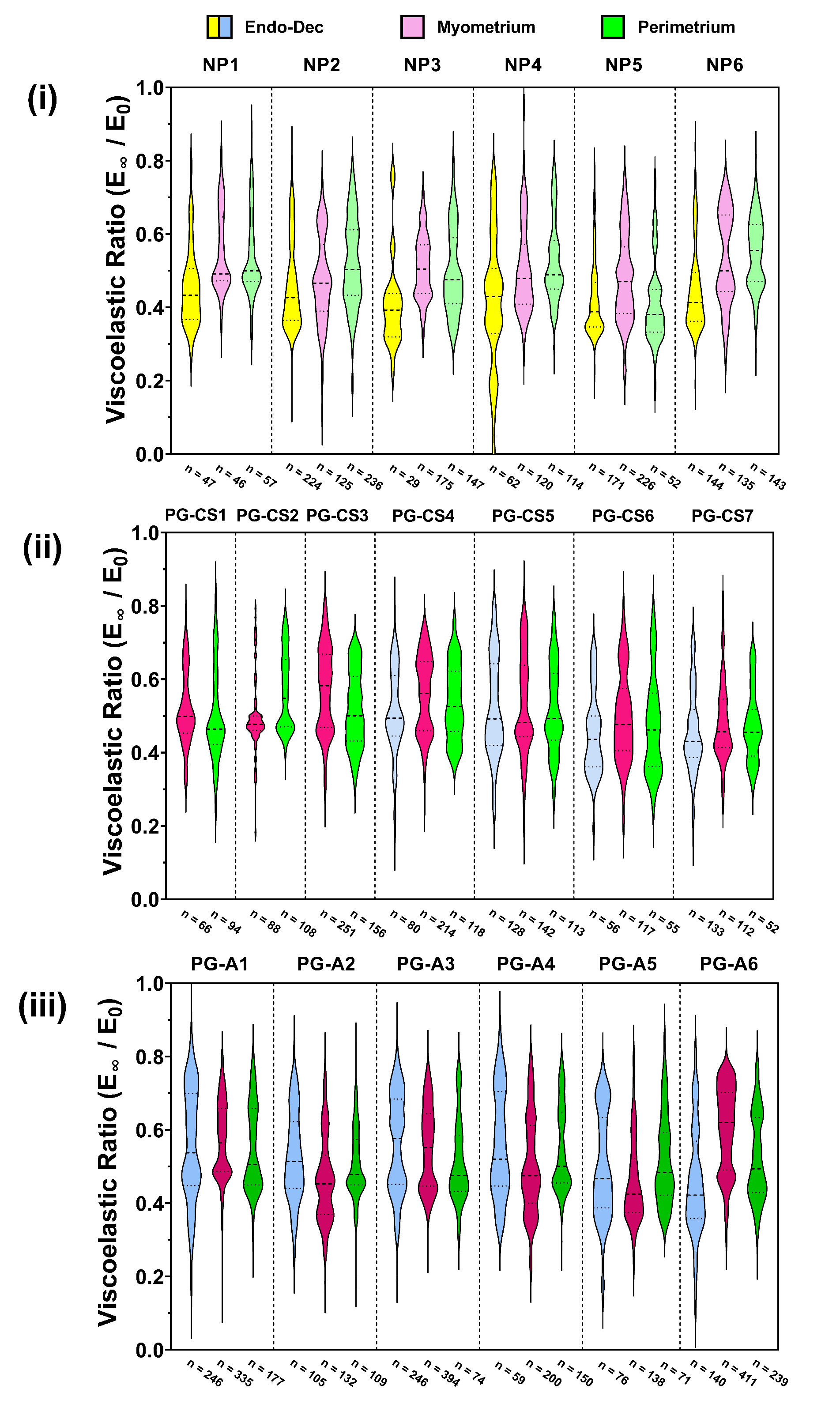

Figure S7. Viscoelastic Ratio Distribution for Individual Patient Uterine Layers. Data are presented as violin plots on a linear scale for each patient group: (i) NP, (ii) PG-CS, (iii) PG-A. ‘n’ denotes technical replicates.

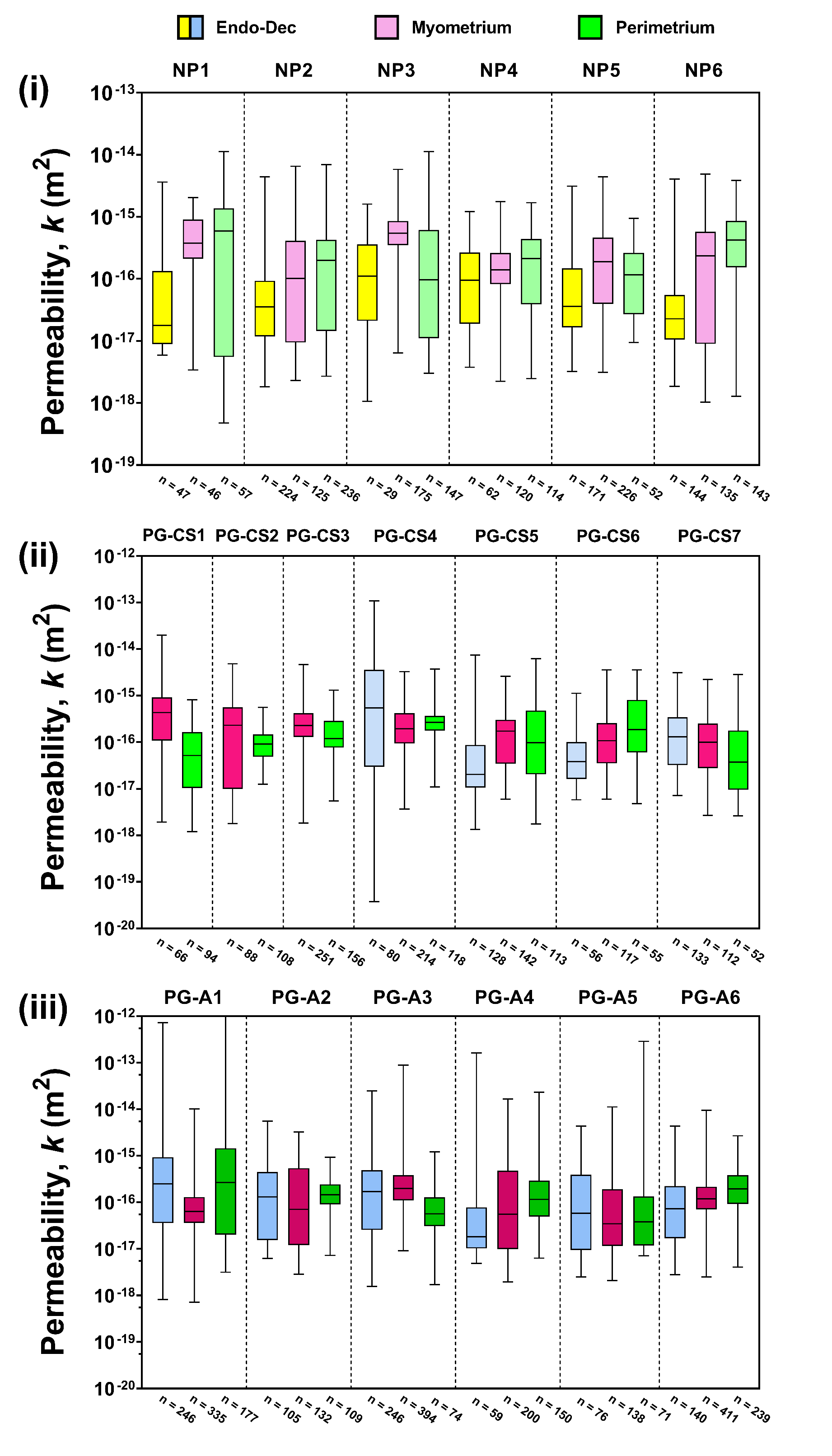

Figure S8. Permeability of Individual Patient Uterine Layers. Data are presented as box and whisker plots on a linear scale for each patient group: (i) NP, (ii) PG-CS, (iii) PG-A. ‘n’ denotes technical replicates.

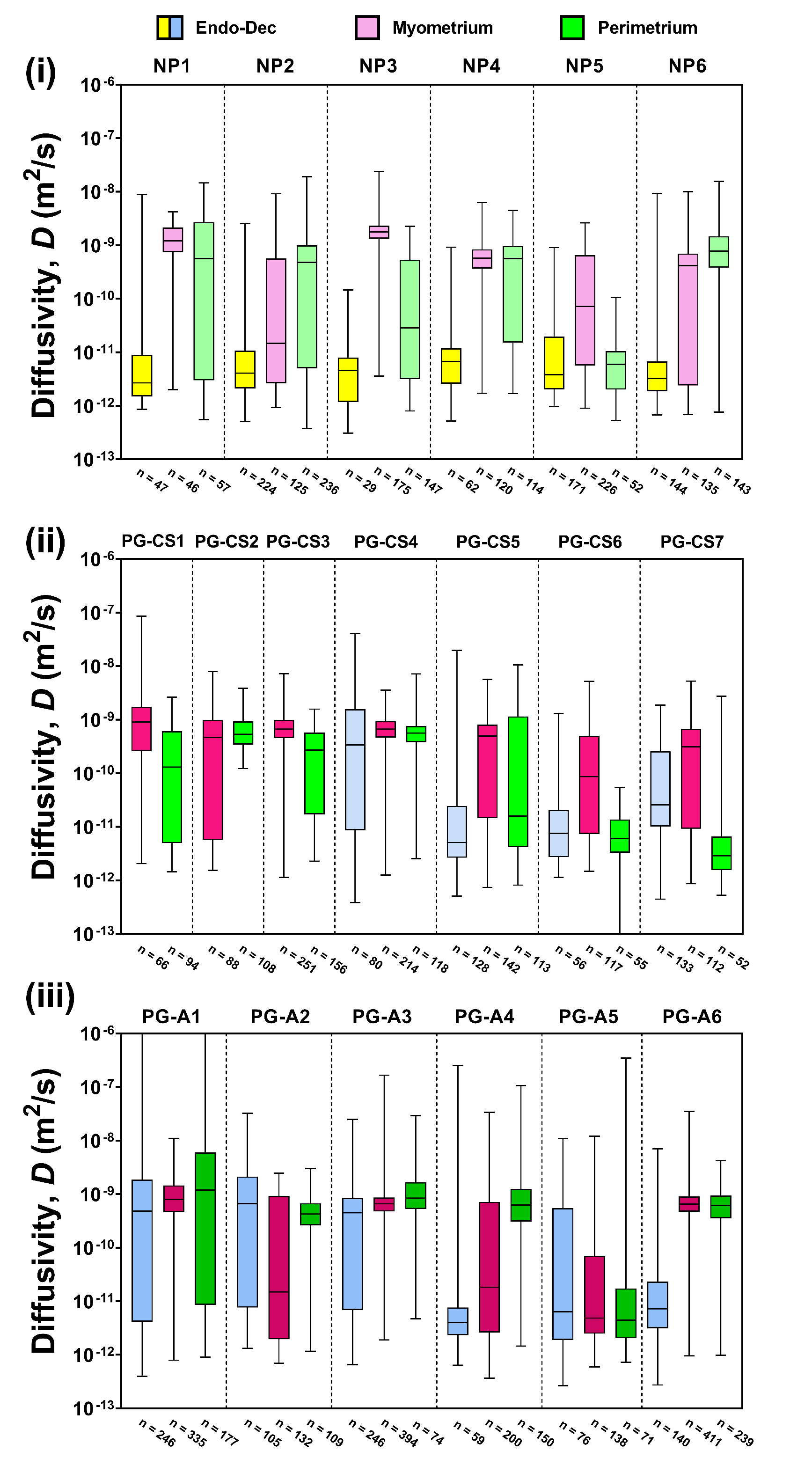

Figure S9. Diffusivity of Individual Patient Uterine Layers. Data are presented as box and whisker plots on a linear scale for each patient group: (i) NP, (ii) PG-CS, (iii) PG-A. ‘n’ denotes technical replicates.

| **Patient ID** | **Age (yrs)** | **Race/ Ethnicity** | **Gest. Age (wks.dys)** | **Placenta Location** | **Gravidity/ Parity** | **Surgery** | **Surgery Indication & Pathological Findings** | **Obstetric History** |
| --- | --- | --- | --- | --- | --- | --- | --- | --- |
| NP1 | 43 | Hispanic | N/A | N/A | 4/3 | Hyst | AUB, PP, Endometriosis, Adenomyosis | Tubal  Ligation |
| NP2 | 36 | Hispanic | N/A | N/A | 3/3 | Hyst | Prolapse  (uterus & vagina), Adenomyosis, Leiomyoma | N/A |
| NP3 | 44 | AA | N/A | N/A | 2/2 | Hyst | PP, Leiomyoma, Adenomyosis | CS x 1 |
| NP4 | 43 | Hispanic | N/A | N/A | 2/2 | Hyst | Leiomyoma | Tubal  Ligation |
| NP5 | 45 | Hispanic | N/A | N/A | 1/0 | Hyst | Leiomyoma | Vtops x 1 |
| NP6 | 47 | White | N/A | N/A | 0/0 | Hyst | Leiomyoma, Menorrhagia, Adenomyosis | N/A |
| PG-CS1 | 42 | White | 39.3 | Post | 3/2 | CS | Repeat CS | CS |
| PG-CS2 | 43 | Hispanic | 37 | Post | 4/3 | CS | Repeat CS | CS x 2 |
| PG-CS3 | 38 | White | 39.1 | Ant | 3/3 | CS | Repeat CS | CS x 2, EA |
| PG-CS4 | 37 | Asian | 38 | Ant | 4/2 | CS | Repeat CS | CS x 1 |
| PG-CS5 | 31 | Hispanic | 39.1 | Ant | 9/5 | CS | Repeat CS | CS x 4 |
| PG-CS6 | 29 | Hispanic | 38.1 | Ant | 2/2 | CS | Repeat CS | CS x 1 |
| PG-CS7 | 33 | Unknown | 39.1 | Ant | 1/1 | CS | Fetal spina bifida Lumbosacral MMC | N/A |
| PG-A1 | 30 | Hispanic | 32 | Post | 4/4 | CS Hyst | Placenta Accreta | CS x 3 |
| PG-A2 | 46 | White | 35.5 | Post | 8/7 | CS Hyst | Placenta Accreta, Placenta Previa | CS x 1,  D&C x 1, VBAC x 2 |
| PG-A3 | 28 | AA | 35.1 | Ant | 5/2 | CS Hyst | Placenta Accreta | CS x 1,  D&C x 1 |
| PG-A4 | 39 | Indian | 35.1 | Ant LLW | 4/3 | CS Hyst | Placenta Accreta | CS x 2 |
| PG-A5 | 33 | Hispanic | 36.3 | Ant + Post | 5/4 | CS Hyst | Placenta Accreta,  Placenta Previa | CS x 2,  SAB x 1 |
| PG-A6 | 38 | White | 36.1 | Ant | 14/12 | CS Hyst | Placenta Accreta | CS x 1,  VBAC x 6 |

Table S1. Detailed Patient Information. [Key] *Placenta Location*: Post = posterior, Ant = anterior, LLW = left lateral wall; *Surgery*: Hyst = hysterectomy, CS = cesarean section; *Surgery Indication*: AUB = abnormal uterine bleeding, PP = pelvic pain; *Obstetric History*: Vtops = voluntary termination of pregnancy, EA = endometrial ablation, D&C = dilation and curettage, SAB = spontaneous abortion, VBAC = vaginal birth after cesarean. *Note*: Parity values include current pregnancy. CS values in obstetric history do not reflect the current pregnancy.

| **Patient ID** | **Estimated Stage of Menstrual Cycle** | **Additional Pathology Notes** |
| --- | --- | --- |
| NP1 | Unknown | Scant endometrium; mostly basalis |
| NP2 | Proliferative | None |
| NP3 | Late Secretory | Histology suggestive of progestin administration |
| NP4 | Secretory | None |
| NP5 | Secretory | Endometrial polyp present |
| NP6 | Unknown | Inactive endometrium |

Table S2. Menstrual Cycle Information – Nonpregnant Patients. Data is estimated from pathology reports.

Data S1. All Nanoindentation Measurements (separate file)

All measurements of stiffness, viscoelastic ratio, permeability, and diffusivity are separated into individual sheets. Each row within a given column represents a discrete indentation point for a given patient’s tissue layer. A final summary data table is included as a separate sheet. The average, standard deviation, and number of indentation points (N) for stiffness, viscoelastic ratio, permeability, and diffusivity are included for each tissue layer corresponding to a different patient.
